# Understanding corticosterone fluctuations and HPA-axis regulation in a mouse model of Spinocerebellar ataxia type 3

**DOI:** 10.64898/2026.01.22.701070

**Authors:** Joana Sofia Correia, Daniela Monteiro-Fernandes, Sara Guerreiro, Bruna Ferreira-Lomba, Daniela Cunha-Garcia, Patrícia Gomes, Andreia Teixeira-Castro, Patrícia Maciel, Sara Duarte-Silva

## Abstract

Spinocerebellar ataxia type 3 (SCA3) or Machado-Joseph Disease (MJD) is a neurodegenerative disease caused by a CAG triplet expansion in the *ATXN3* gene, primarily characterized by motor impairments. However, SCA3/MJD also includes mood-related comorbidities that affect both patients and their caregivers. Current treatments focus on symptom management and supportive care, as no disease-modifying therapies are available. Previously, we have demonstrated decreased glucocorticoid receptor (GR) expression in post-mortem SCA3 human and mouse brains and elevated peripheral corticosterone (CORT) levels in SCA3 mice at late disease stages. Impaired GR signaling is typically associated with hypothalamic-pituitary-adrenal (HPA) axis dysfunction, common in stress-related psychiatric diseases. Our study aimed to dissect the HPA-axis (dys)function and the effect of stress exposure on SCA3/MJD progression. Using the CMVMJD135 mouse model, we evaluated HPA-axis regulation in SCA3/MJD by measuring CORT levels throughout disease progression (6 to 34 weeks of age) under basal conditions and after acute stress. At week 35, these mice underwent a dexamethasone injection to challenge the HPA-axis, and the CORT levels were measured at different timepoints to evaluate the axis response. Additionally, we applied a 6-week chronic unpredictable stress (CUS) protocol in another cohort of mice starting at an early symptomatic stage to assess stress effects on the progression of motor impairments. Our findings indicate that serum CORT levels in SCA3 mice begin to rise between 26 to 30 weeks of age, with no impairment in the physiological response to acute stress. SCA3 mice were also able to normalize CORT levels after dexamethasone challenge, suggesting normal HPA-axis function. While CUS exposure had a transient negative impact on the motor phenotype, this effect did not persist throughout disease progression. In conclusion, stressful events, either acute or chronic, do not seem to be major determinants of disease severity in SCA3 mice.

## 1. Introduction

Spinocerebellar ataxia type 3 (SCA3), also known as Machado-Joseph disease (MJD), is an autosomal dominant inherited polyglutamine disease caused by a CAG expansion in the *ATXN3* gene ^1^. This expansion alters the ataxin-3 protein structure, causing abnormally long glutamine tracts, and subsequently promoting protein misfolding and aggregation throughout the brain ^2, 3^. The nuclear accumulation of protein aggregates in neurons combined with progressive disruption of normal cellular functions, impaired protein homeostasis, and increased oxidative stress processes, together contribute to a vicious cycle of proteotoxicity in cells that ultimately leads to progressive neuropathology (reviewed in ^4, 5^). Clinically, SCA3/MJD is represented by progressive ataxia, ophthalmoplegia, amyotrophy, dystonia, and/or spasticity ^2, 6^. Although efforts have been made towards understanding the processes of neurodegeneration behind this disease, no effective treatment is yet available. Importantly, reports of anxiety and depressive symptoms are common for SCA3/MJD patients. In a comprehensive study, Lo *et. al*. (2016), reported that depression is a common comorbidity in Spinocerebellar Ataxia (SCA) group of diseases, with a prevalence rate of 26% ^7^. While the baseline prevalence of depression is similar across various SCA types, suicidal ideation is notably higher in SCA3/MJD, affecting about 65% of the patients from this cohort. Depressive symptoms were correlated with SARA (Scale for the Assessment and Rating of Ataxia) scores but did not show significant progression over a two-year period, nor it was detected a direct relationship with increasing number of pathological CAG repeats ^7^.

Over the past years, our team has been dedicated to the development of therapeutic interventions and the identification of therapeutic targets for SCA3/MJD ^8-13^. Among effective treatments, the antidepressant citalopram was observed to successfully ameliorate motor phenotype of SCA3/MJD animals, demonstrating higher neuroprotective effects when administered prior to disease onset ^9^, but still being beneficial when administered upon disease installation ^9, 10^. Recently, we showed a significant reduction of GR protein levels in the brainstem of SCA3 mice, which were also observed in the pons of postmortem SCA3 samples. Additionally, SCA3/MJD patients also exhibited a reduction in GR in the blood ^11^. To better understand the reduced levels of GR in the context of SCA3/MJD, we proved that GR physically interacts with ATXN3 and that levels of ubiquitylated GR were increased in the brain of the SCA3/MJD mouse ^11^. These results strongly point to an involvement of ATXN3 in the degradation process of GR, an hypothesis that requires further studies. The obtained results, however, put forward the hypothesis of a relevant role for the HPA-axis in the context of SCA3/MJD. Despite exhibiting normal daily fluctuations in corticosterone (CORT) levels at a late stage of the disease, the baseline CORT levels of these mice were elevated compared to healthy control animals. Again, this suggests a potential dysfunction in the GR-dependent signaling pathways in the context of SCA3/MJD.

The temporal activation of the HPA axis is influenced by the duration and type of stressor. The acute stress response effectively triggers the HPA axis activation, which is subsequently regulated by feedback mechanisms that help to terminate the response once the stressor has ended. Typically, the HPA response is initiated by a surge of adrenocorticotropic hormone (ACTH), which begins within minutes and lasts for a brief period, which is contingent on the length and intensity of the stimulus as well as feedback processes. The glucocorticoid response, in contrast, has a delayed onset due to the need for new glucocorticoid production in the adrenal glands and persists for a considerably longer period. Thus, the duration of this response is determined by both active feedback signaling and passive processes, such as glucocorticoid degradation. Impaired GR signaling is regarded as a key mechanism in the pathogenesis of stress-related psychiatric diseases, namely, major depressive disorder, which studies point out to a decrease in peripheral GR expression ^14-16^. Given the pivotal role of GR mechanisms in the response of the HPA axis to stress stimuli ^17-20^, and our previous results ^11^, it was of our interest to ascertain whether SCA3 mice exhibited susceptibility to stress exposure. To dissect HPA-axis regulation in the CMVMJD135 transgenic mice, a longitudinal quantification of serum CORT was performed at multiple timepoints throughout disease progression (from 6 weeks to 34 weeks of age) under basal conditions and following an acute stressor. In addition, an unpredictable chronic stress (CUS) protocol was conducted in this animal model at young-adult ages (from 6 to 12 weeks), to mimic a possible stress inference on SCA3/MJD patients upon prior knowledge of carrying such devastating disease. Our results demonstrated an elevation of basal CORT levels from 26 weeks of age onwards, an age where motor symptoms and neuropathology are fully established. Despite this elevation, the fluctuations in CORT levels in response to acute stress were similar between SCA3 mice and WT animals, indicating a normal stress response. Furthermore, the dexamethasone suppression test confirmed that the HPA axis in SCA3 mice remains functional, as they were able to restore CORT levels after axis suppression. Notably, when exposed to chronic stress, SCA3 mice displayed only mild and temporary worsening of motor symptoms, suggesting resilience to prolonged stress exposure.

## 2. Methods

### 2.1 Animal experimentation ethical statement

All procedures were conducted in accordance with European regulations (European Union Directive 86/609/EEC), transposed to the Portuguese law according to the paragraph a) of article 31º of Decree-Law 113/2013 of 7 August with the alterations introduced by the Decree-Law 1/2019 of January 10. Animal facilities and experimenters were certified by the Portuguese regulatory entity (Direcção Geral de Alimentação e Veterinária -DGAV). All protocols were approved by the Animal Ethics Committee of the Life and Health Sciences Research Institute, University of Minho and by the DGAV (reference 020317).

### 2.2 Animal model and living conditions

Both female (F) and male (M) CMVMJD135 (C57BL/6J background, hereon referred as SCA3 mice for simplicity) transgenic mice were used in the study. At 4 weeks-old, SCA3 mice and wild-type (WT) littermates were assigned by sex and housed at weaning in groups of 6 animals in filter-topped polysulfone cages 267 × 207 × 140 mm (370 cm2 floor area) (Tecniplast, Buguggiate, Italy), with corncob bedding (Scobis Due, Mucedola SRL, Settimo Milanese, Italy) and nesting material in a conventional animal facility. SCA3 mice were mated with WT animals, and their progeny were genotyped at weaning using PCR. Genotyping procedures were previously described ^21^. Mean CAG repeat size is reported in the protocol methods.

All animals were maintained under standard laboratory conditions: an artificial 12h light/dark cycle (lights on from 8 a.m. to 8 p.m.), with a room temperature of 21±1°C and a relative humidity of 50–60%). Mice were given a standard diet (4RF25 during the gestation and postnatal periods, and 4RF21 after weaning) (Mucedola SRL, Settimo Milanese, Italy) and water *ad libitum*. Health monitoring was performed according to FELASA guidelines, confirming the Specified Pathogens status of sentinel animals maintained in the same animal room. Humane endpoints for the experiment were defined (20% reduction of the body weight, inability to reach food and water, presence of wounds in the body, dehydration) but not needed in practice as the study period was conceived to include ages at which animals do not reach these endpoints.

### 2.3 Corticosterone measures

Two different groups of mice were used for CORT measures: (1) a first group of mice were subjected to independent episodes of acute stress at 18, 26 and 34 weeks of age, and blood was collected prior to the stressor, and 15 and 90 minutes after stress exposure (Figure1A-B; WT: *n* = 5, 5F; SCA3: *n* = 9, 9F; 142 CAGs ± 1.2 (mean ± SEM); 136–146 (min– max)_CAG_); (2) in a second group of animals serum CORT quantification was performed at several timepoints during disease progression (from 6 to 34 weeks of age) at basal conditions (8 AM and 8 PM; Figure 2A; WT: *n* = 10, 5F/5M; SCA3: *n* = 10, 6F/4M; 142 CAGs ± 0.9 (mean ± SEM); 139–148 (min–max)_CAG_), accordingly to the CORT cycle.

**Figure 1.**
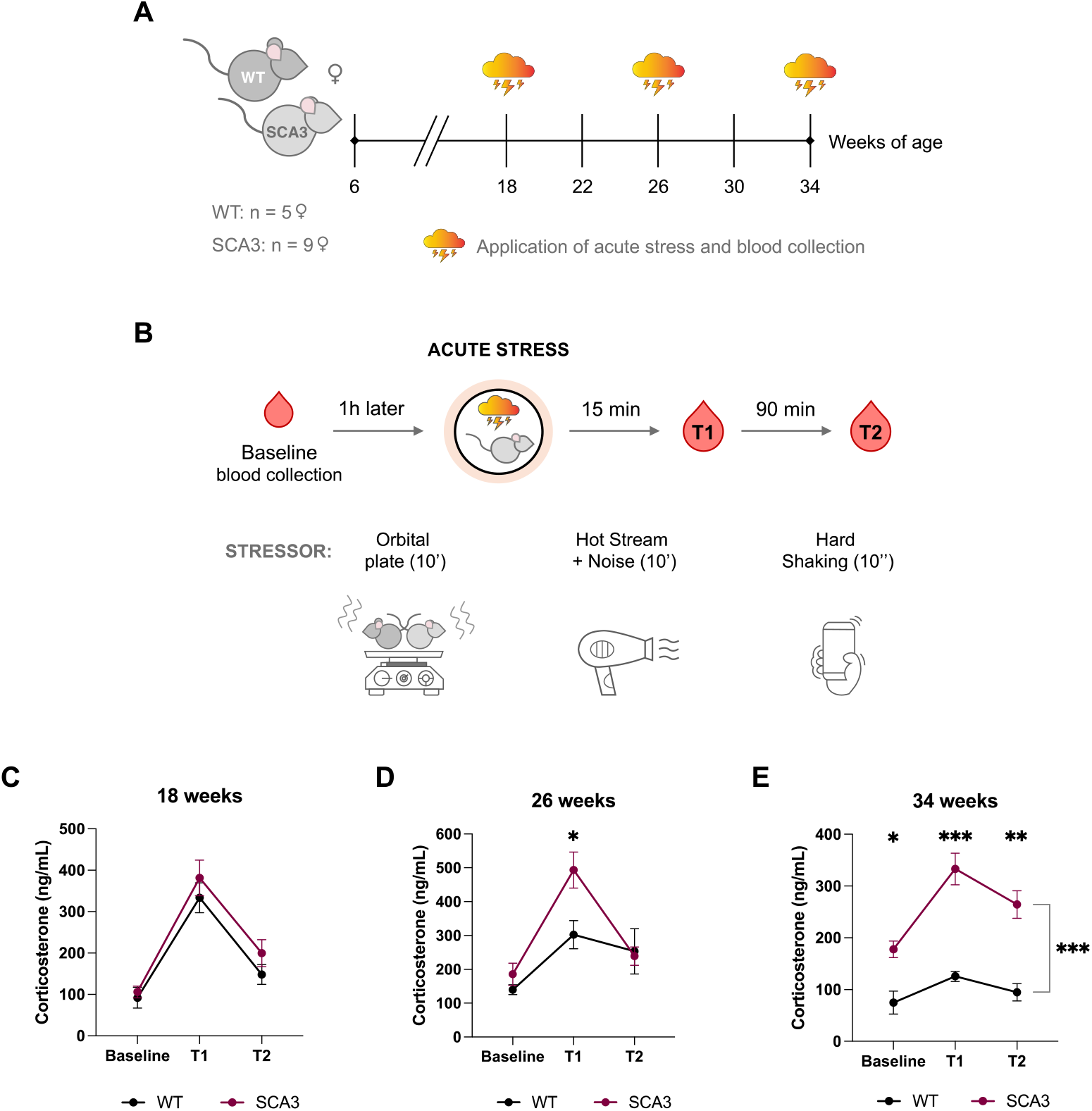
SCA3 mice could rebalance the blood CORT levels after an acute stress event. (A) Timeline representation of the acute stress experiment along with disease progression. (B) Schematic representation of the acute stress protocol; different stressors were applied to avoid re-exposure biases. (C) At 18 weeks of age SCA3 mice present a similar response of corticosterone (CORT) variation to their wild-type (WT) counterparts before and after the application of the acute stressor (10 minutes in orbital plate rotating at 140 RPM). (D) SCA3 mice with 26 weeks of age could restore the CORT levels in blood after the exposure to 10 minutes of hot air and noise. At a late stage of the disease, 34 weeks-old SCA3 mice present increased CORT levels at the baseline, when compared to WT animals. This elevation in CORT was observed in all timepoints measured at this age, nonetheless SCA3 mice show a rebalance response 90 minutes after the 10 seconds of hard shaking acute stressor (E). WT – wild-type; SCA3 – CMVMJD135 mouse; Baseline – 1 hour prior to stressor; T1 – timepoint of 15 minutes after acute stressor; T2 – timepoint of 90 minutes after acute stressor. A mixed-design repeated measures ANOVA was applied in statistical analyses of continuous variables with normal distribution (C–E). Data is represented as the mean ± SEM. * p < 0.05; ** p < 0.01; *** p < 0.001.

**Figure 2.**
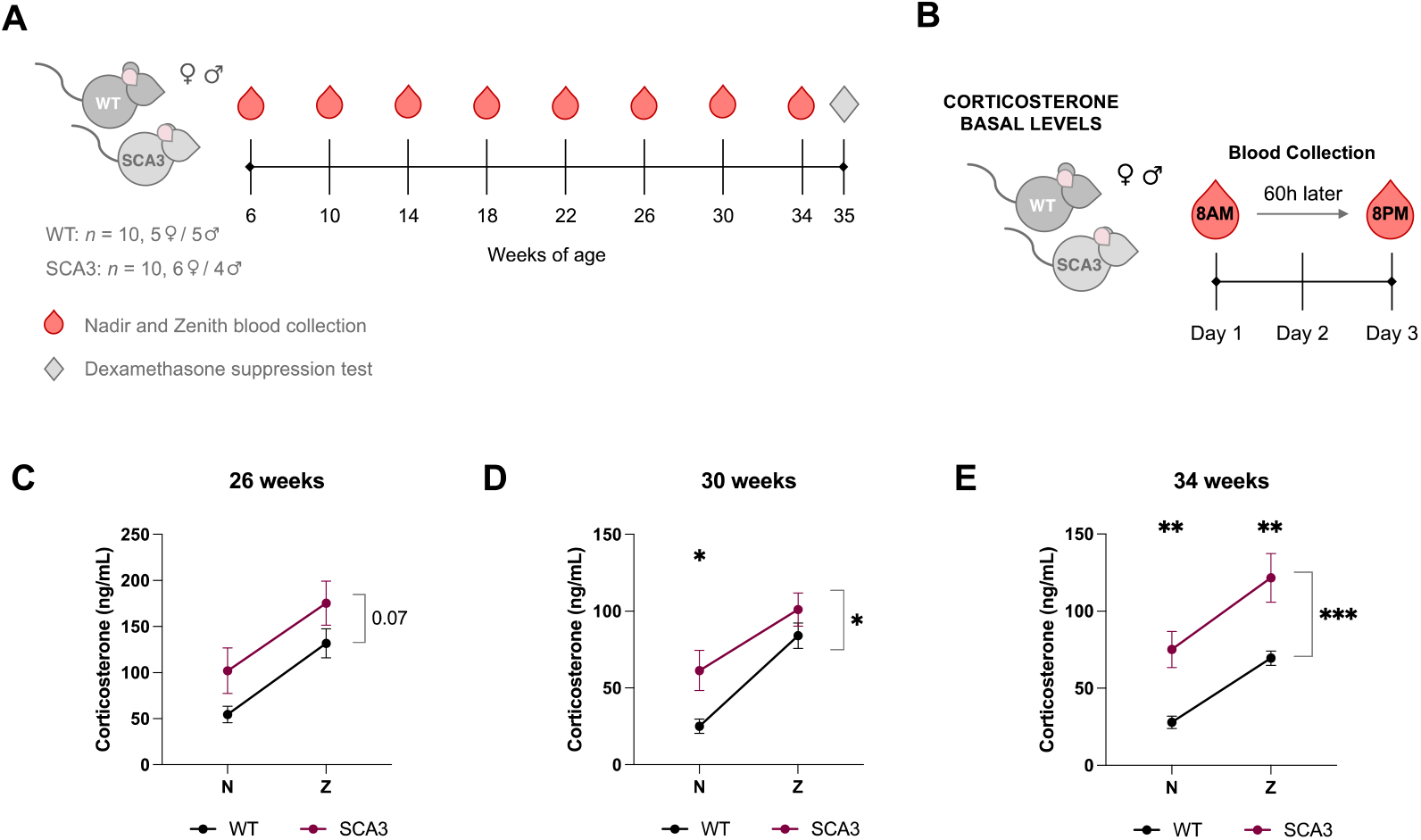
There is a CORT elevation in the blood of SCA3 mice as disease progresses. (A) A longitudinal analysis was conducted in the SCA3/MJD mouse (SCA3) to evaluate corticosterone (CORT) levels as disease progresses. (B) Blood collection protocol for the assessment of corticosterone (CORT) levels at the nadir (8AM) and zenith (8PM), at late stages of SCA3/MJD. (C-E) An elevation of CORT levels was observed throughout disease progression. Starting with a tendency to increase at 26 weeks of age (C) and reaching significance at 30 weeks of age (D); (E) CORT levels became elevated in both timepoints at 34 weeks of age. WT – wild-type; SCA3 – CMVMJD135 mouse; N – Nadir; Z – Zenith. Continuous variables with normal distributions were analyzed using mixed-design repeated measures ANOVA (C–E). Data is represented as the mean ± SEM. * represents statistical differences for genotype; # represents statistical differences between sex for the WT animals; * p < 0.05; ** p < 0.01; *** p < 0.001; ## p < 0.01.

For both cases, at each timepoint, about 20–30 µL of blood were collected to a single-use polypropylene tube by tail snip. After blood collection, samples were centrifuged for 20 minutes at 10 000 RPM to collect the serum to a new tube and stored at -80ºC. To read the CORT levels an ELISA assay was performed using either the Enzo-kit (1/40 dilution, ADI-90-097, Enzo Life Sciences, Inc., NY, USA) for basal CORT analyses or the Abcam ELISA kit (ab108821, Abcam Limited, CB2 0AX, UK) for the acute stress experiment.

### 2.4 Acute Stress

At week 18, 26, and 34, mice were submitted to acute stress and blood was collected at baseline conditions one hour prior the stressor application, followed by blood collection 15 and 90 minutes after the stress exposure. For ethical reasons, we used a different ELISA kit (ab108821, Abcam Limited, CB2 0AX, UK) because it requires a smaller amount of serum (2 µL; 1/70 dilution). This allowed us to minimize the volume of blood collected at each time point, ensuring the animals’ welfare was prioritized.

To avoid desensitization, different types of acute stressors were applied to the same mice at three different timepoints separated by 2 months each. The stressors applied included: 10 minutes of 140 RPM shaking in an orbital plate, 10 minutes of hot stream (hairdryer), and 10 seconds of manual hard shacking.

### 2.5 Dexamethasone Suppression Test

Using the same animal cohort from basal CORT measures, at 35 weeks of age, after the last timepoint blood collection, the dexamethasone suppression test (DST) was performed accordingly to previous described protocols ^22^. Briefly, blood was collected 1 hour prior receiving a single intraperitoneal injection of dexamethasone (0.1 mg/Kg in 0.9 % saline; Fortecortin® Inject 4 mg/mL, Merck KGaA, Darmstadt, Germany) in the morning period, to establish baseline levels. 6 hours after dexamethasone injection, the animals were euthanized, and blood was collected for CORT analysis. CORT levels were measured using the Enzo-kit (1/40 dilution; ADI-90-097, Enzo Life Sciences, Inc., NY, USA).

### 2.6 Chronic unpredictable stress protocol

Both male and female WT and SCA3 mice with 6 weeks of age were exposed to a chronic unpredictable stress (CUS) paradigm for 6 weeks ^23, 24^. Different types of stressors were applied unpredictably, one period of stress per day, consisting of exposure to incrementing times of either restraint (1 – 5 hours), vibrating platform (15 minutes – 5 hours), overcrowding (1.5 – 5 hours), or hot air stream (15 – 20 minutes) on consecutive days. To monitor the efficacy of CUS, body weights were measured weekly, and blood was collected at the fourth week of the protocol and assayed for CORT levels. Additionally, animals performed behavioral tests both at the 4^th^ and 6^th^ weeks of the CUS protocol to evaluate anxiety and depressed-like phenotypes (coping behavior) induced by stress exposure. During the CUS protocol, both WT and SCA3 control littermates were left undisturbed in their home cages and handled by the same experimenter for 20 minutes to a total of 1 hour in one period per week. 10 males of both genotypes were lost during 5 hours of overcrowding in the 6^th^ week of CUS application. Also, the 5 hours of stress exposure were proved to be too aggressive for these mice. This procedure was stopped immediately, and the periods of exposure were limited to maximum of 4 hours. Four groups of mice were than established in the motor analyses of this experiment: WT (control group, *n* = 17, 11F/6M); SCA3 (control group, *n* = 16, 8F/8M; 140 CAGs ± 0.7 (mean ± SEM); 137–147 (min–max)_CAG_); WT-CUS (stressed group, *n* = 11, 9F/2M); SCA3-CUS (stressed group, *n* = 11, 9F/2M; 142 CAGs ± 1.0 (mean ± SEM); 138–147 (min–max)_CAG_).

After CUS exposure the motor behavior of these animals was evaluated in a longitudinal assessment. At week 34, the animals were deeply anesthetized with a mixture of ketamine hydrochloride (150 mg/kg) plus medetomidine (0.3 mg/kg) and euthanized by exsanguination perfusion with saline or PFA 4%. Tissue samples were stored at -80ºC or conserved in PFA 4% at 4ºC, accordingly.

### 2.7 Mouse behavior analysis

All animals included in this study (*n* = 10–17 per genotype) were assessed for behavior. All procedures are explained in the following sections.

#### 2.7.1 Mouse motor phenotype

All mice were weighed from 5 to 34 weeks of age every two weeks. To control for motor deficits, animals performed several behavioral paradigms, including the motor swimming and beam balance tests, the hanging wire grid, spontaneous horizontal activity, and gait quality assessment. All these measures were previously used and validated in this mouse model of SCA3 ^8, 9, 13, 25^.

#### 2.7.2 Behavioral tests to evaluate CUS efficacy

##### Forced Swimming Test (FST)

Alterations in coping behavior caused by chronic stress exposure is measured by learned helplessness paradigms such as the FST ^26^. For that, mice were placed in glass cylinders filled with water (23 °C; 50 cm high) for 6 minutes (min), and the test video-recorded and scored manually for the last 4 min of the trial. An increase in immobility time was taken as a measure of low coping behavior at the 6^th^ week of the CUS paradigm (WT (*n* = 17, 11F/6M); SCA3 (*n* = 16, 8F/8M; 140 CAGs ± 0.7 (mean ± SEM); 137– 147 (min–max)_CAG_); WT-CUS (*n* = 11, 9F/2M); SCA3-CUS (*n* = 10, 8F/2M; 142 CAGs ± 1.0 (mean ± SEM); 138–147 (min–max)_CAG_).

##### Light-Dark Box (LDB)

The LDB test was performed at the 6^th^ week of CUS to access anxiety levels in stressed mice. The apparatus for this test was the OF arena divided in half using a black acrylic box placed at the right of the arena. In this test the animals are allowed to explore between an illuminated area versus a dark area/box for a maximum of 10 min ^27^. The trajectory of each mouse was detected automatically through infrared beams and using the MedAssociates Inc. (St Albans, VT, USA) software. Lower percentage of time spent exploring the light area was counted as anxious behavior.

### 2.8 Statistics

In this study, a single animal served as the experimental unit. The sample size was determined using power analysis and estimates as previously described ^8, 9^. Continuous variables with normal distributions (Shapiro–Wilk test p > 0.05) were analyzed using Student’s t test, repeated measures ANOVA (factors were timepoint and group), two-way ANOVA (factors were stress and genotype; Tuckey’s post hoc test was used for multiple comparisons). All continuous data are shown as the as mean ± standard error of the mean (SEM). Whenever a continuous variable did not present a normal distribution the nonparametric Mann–Whitney U test was applied. Whenever ANOVA variables fail the Levene’s test for Equality of variances, the Dunnet’s post hoc test was considered in the multiple comparisons. Discrete and categorical variables were analyzed using the nonparametric Mann Whitney U test (when two groups) and the Kruskal-Wallis H (more than two groups). Linear regression model was applied to determine correlation between variables and *Pearson’s r* and *r* ^*2*^ both considered as correlation coefficients. Numbers above 1.5*IQR were considered outliers and excluded from the analyses. All statistical analysis was performed using SPSS 29.0 (IBM Corp., Armonk, NY, USA). Significance was accepted for a critical value of p < 0.05.

## 3. Results

A longitudinal analysis was performed in the CMVMJD135 mouse model, to assess CORT levels after an acute stress stimulus (Figure 1A) and at basal levels (Figure 2A) throughout disease progression. Different acute stressors were applied to avoid habituation (Figure 1C). SCA3 animals used for both experiments had a low variation of the CAG triplet number, resulting in a similar disease severity between experimental groups: basal group, 142 ± 0.9 (CAG_n_, mean ± SEM); acute stress group, 142 ± 1.2 (CAG_n_, mean ± SEM; Supplementary Figure S1A).

### 3.1 SCA3 mice rebalanced CORT levels after an acute stress event

Starting at a mid-stage of the disease ^8, 9^, 18 weeks-old SCA3 mice did not present an alteration of baseline CORT levels nor of HPA-axis stress-dependent response when compared to WT counterparts, herein translated as CORT variations in serum (Figure 1C). Although the baseline levels of both groups still not reach the statistical difference at 26 weeks-old, SCA3 mice clearly had higher CORT response 15’ after stressor application (Figure 1D). At this age, CORT levels were well-restored after 90’ of rest. However, at 34 weeks of age, a late-stage of the disease, SCA3 mice presented higher levels of CORT in serum (Figure 1E, Baseline), as previously observed ^11^. Despite this basal increase, SCA3 mice could restore CORT to baseline levels at the second timepoint (90’ of rest, Figure 1E, T2; SCA3 Baseline x T2, F_(1, 10)_ = 48.68; p = 0.1036), indicating proper HPA-axis mediation of the stress response.

### 3.2 SCA3 mice exhibited increased CORT levels at mid to late-stage of the disease

To map the onset of CORT levels elevation in SCA3 mice, blood was collected both at nadir (8AM) and zenith (8PM) timepoints in different disease stages (Figure 2A). The collection was separated for 60 hours to respect animals’ welfare (Figure 2B). Like what was observed in the acute stress experiment, 26 weeks-old SCA3 mice did not show statistical difference to their WT littermates in CORT levels at neither nadir nor zenith timepoints, despite a clear trend towards increased CORT levels in SCA3 animals (p=0.07) (Figure 2C). However, this alteration achieved its statistical difference at 30 weeks of age, particularly at nadir CORT levels (Figure 2D). As expected from previous results ^11^, increased levels of peripheral CORT were observed in SCA3 mice at 34 weeks of age (Figure 2E). Because no statistical differences were observed at 26 weeks of age, the blood collected at previous timepoints (see timeline in Figure 2A) were not analyzed.

### 3.3 SCA3 mice showed typical CORT fluctuations in response to dexamethasone-induced HPA-axis challenge

At 35-weeks of age animals were challenged with a single shot of dexamethasone to suppress HPA-axis CORT response (Figure 3A). In this test we could observe a successful action of dexamethasone to compete for corticosterone receptors at the hypothalamus creating a negative feedback response of the HPA-axis (see Figure S1B for HPA-axis elucidation). This effect was also observed in SCA3 mice 6 hours after dexamethasone administration (Figure 2B). Again, we confirmed the increased CORT levels in 35 weeks-old SCA3 mice one-hour prior injection (Figure 2B, Baseline timepoint). The following week after dexamethasone exposure, animals were euthanized. Interestingly, SCA3 female mice had significantly lighter adrenal glands in comparison to WT (Figure 2C). A negative correlation was observed between baseline CORT levels in 35 weeks-old SCA3 mice and adrenal glands weight, but no difference regarding animals’ sex (Figure 2D).

**Figure 3.**
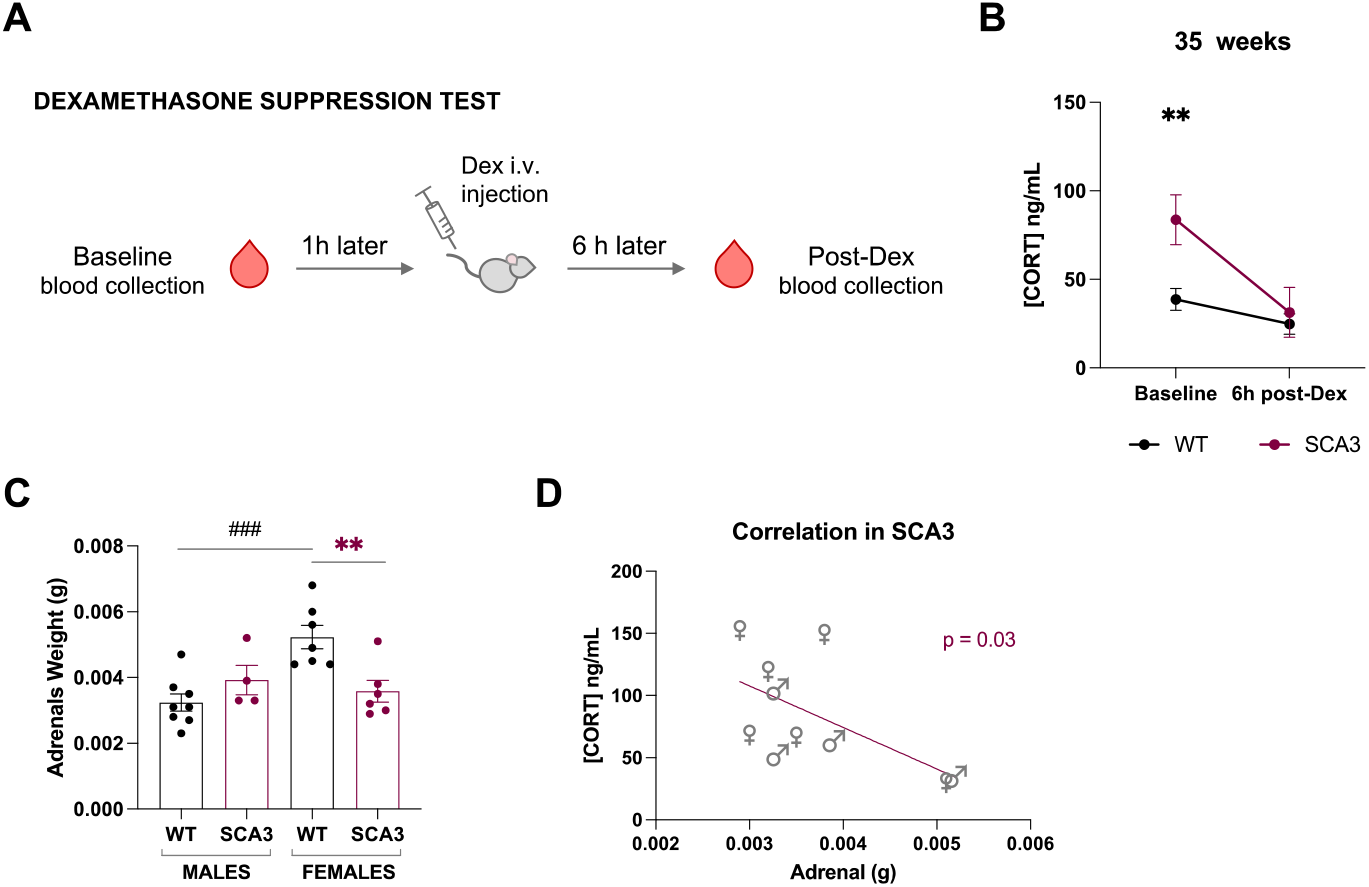
HPA-axis response to external stimuli is not affected by the elevation of CORT in the blood of SCA3 mice as disease progresses. At 35 weeks of age, SCA3 mice were submitted to a (A) protocol for suppressing the HPA-axis response by a single intravenous injection of dexamethasone (Dex). (B) At this age we confirmed that there is an elevation of CORT levels in the blood of SCA3 mice, which were reduced by Dex 6 hours post-injection, therefore suppressing the HPA-axis responses. The baseline corresponds to blood collection 1 hour before Dex injection. (C) SCA3 female mice had significantly lighter adrenal glands in comparison to WT, and (D) a negative correlation was observed between CORT levels in 35 weeks-old SCA3 mice and the respective adrenal glands weight, but no sex differences were found. WT – wild-type; SCA3 – CMVMJD135 mouse. Continuous variables with normal distributions were analyzed using mixed-design repeated measures ANOVA (B), and two-way ANOVA (C). Linear regression model was applied to determine correlation between variables (D). Data is represented as the mean ± SEM. * represents statistical differences for genotype; # represents statistical differences between sex for the WT animals; * p < 0.05; ** p < 0.01; *** p < 0.001; ## p < 0.01.

### 3.4 Chronic stress exposure did not significantly aggravate the prospects of SCA3/MJD progression

A longitudinal evaluation of motor performance was conducted to determine the effects of a 6-week period of stress exposure on SCA3/MJD progression in mice (short- and long-term analysis post-stress; Figure 4A). CAG triplet mean was balanced in SCA3 mice control and stressed groups to avert differences in disease severity (Figure 4B). At week 4 of the CUS protocol blood was collected to check for stress influence in daily CORT cycle (Figure 4C). The expected CORT peak at zenith was observed for both genotypes in basal conditions (WT and SCA3, Figure 4C). A successful effect of CUS protocol in the circadian rhythm of stressed WT mice was achieved as no difference between nadir and zenith timepoints were observed for this group (WT-CUS, Figure 4C). Unexpectedly, this was not the case for SCA3 mice (SCA3-CUS, Figure 4C). In addition, a decrease in weight gain was observed in stressed groups, for both WT and SCA3 mice (Figure 4D), proving protocol efficacy after 6 weeks of exposure. To assess CUS protocol efficacy on inducing learned helplessness, a FST was applied at the end of the 6 weeks period (Figure 4E). A deficit in this coping behavior was observed in both WT-CUS and SCA3-CUS groups (comparing to their non-stressed genotypes) suggesting a depressive-like effect in stressed mice given by their increased immobility time (Figure 4E). An anxious phenotype was observed in the WT-CUS group (Supplementary Figure S2A). At this point, we can conclude that the CUS protocol had the expected phenotypic impact on mice.

**Figure 4.**
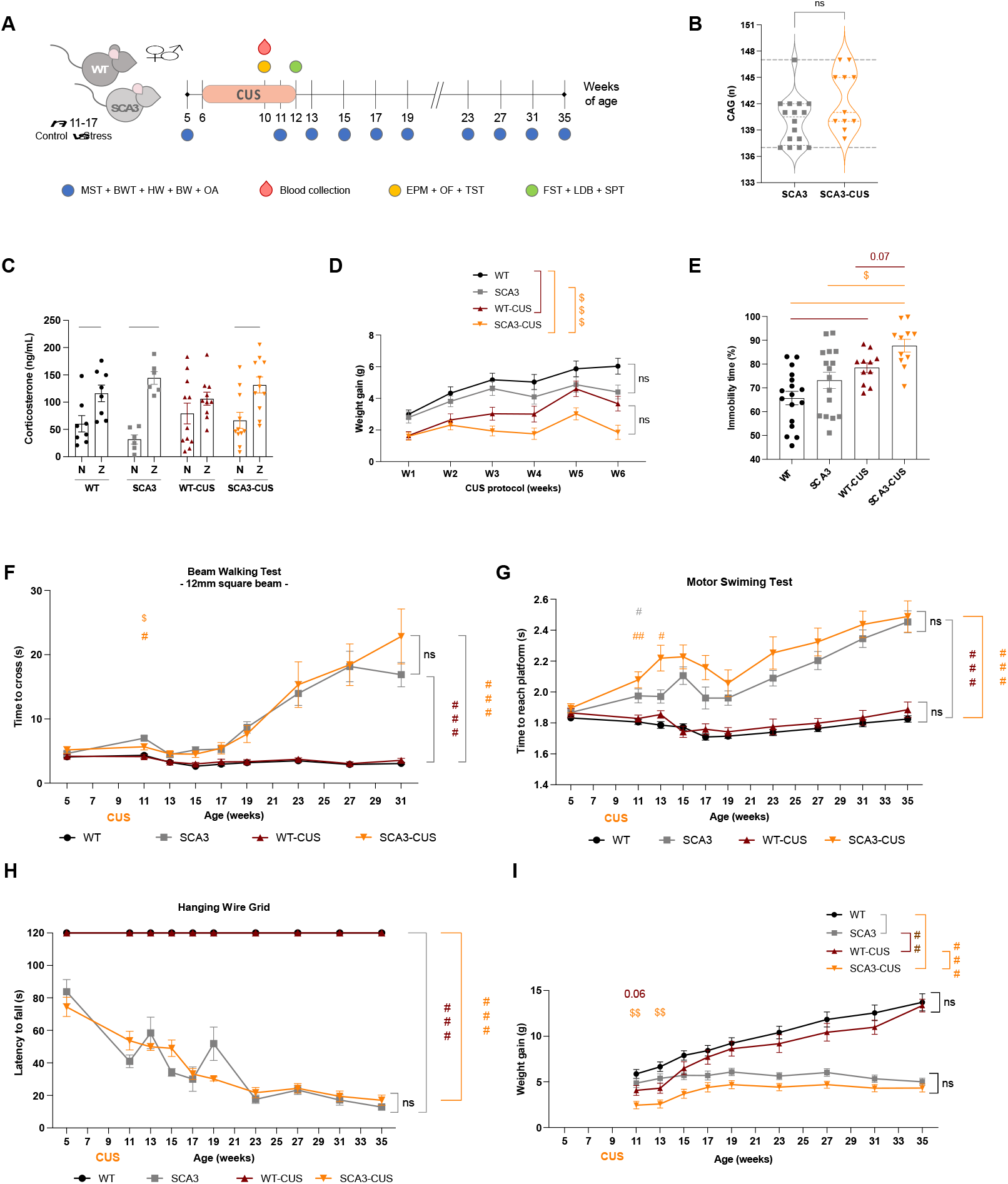
Chronic stress at early age had no major impact in SCA3 mice motor performance. (A) Chronic unpredictable stress (CUS) was applied in 6 weeks-old mice of both sexes and genotypes for 6 consecutive weeks, and their motor performance was measured prior to CUS initiation (at 6 weeks of age), at the end of CUS protocol (at 12 weeks of age) and in several timepoints (blue circles) throughout disease progression until week 35. (B) The CAG repeat expansion of both control and stressed animals were determined and no statistical difference was found between groups of SCA3 mice. (C) After 4 weeks of CUS protocol, a variation of corticosterone (CORT) nadir (8 AM) and zenith (8PM) levels was observed in stressed WT animals, validating CUS protocol. (D) Lower weight gain was observed between WT-CUS animals and the non-stressed WT group, during CUS exposure. To determine influences of CUS at the behavioral level of mice (E) the forced-swimming test was applied, with which we also observe an increased immobility state for the WT-CUS group compared to WT. SCA3-CUS mice showed better balance assessed in the (F) 12mm square beam at 11 weeks of age, but that effect did not sustain over time. No impact of stress was observed for both genotypes in motor coordination in the (G) motor swimming test or muscular strength measured by (H) the hanging wire grid test. Comparing to non-stressed SCA3 mice, SCA3-CUS group exhibited significantly lower (I) weight gain between 11 and 13 weeks of age but losing those effects over time. Genotype comparisons were analyzed using a nonparametric Mann-Whitney U test (B). Repeated measures ANOVA (C-D, F-I) and two-way ANOVA (E) tests were applied in statistical analyses of continuous variables with normal distribution. WT – wild-type mice; WT-CUS – stressed wild-type mice; SCA3 – CMVMJD135 mice; SCA3-CUS – stressed CMVMJD135 mice. Data is represented as the mean ± SEM. * represents statistical differences to non-stressed WT group; # represents statistical differences to WT-CUS group; $ represents statistical differences between non-stressed SCA3 and SCA3-CUS groups; *, $, # p < 0.05; **, $$, ## p < 0.01; ***, $$$, ### p < 0.001; ns: not significant.

To properly infer both the immediate and long-term effects of stress exposure, animals were evaluated for their baseline motor performance at 5 weeks of age, before stress exposure. At this age, all experimental groups showed similar motor performance (Figure 4F-H). Pertinent for comparisons, stress had no impact on WT mice motor performance along time on both balance and coordination measures (Figure 4F-G). Intriguingly, at the last week of CUS protocol, SCA3 mice exposed to stress showed better balance when compared to non-stressed controls, however, this effect did not sustain over time (Figure 4F, S2B-D). Although not statistically significant, stressed-SCA3 mice presented a worst swimming performance throughout disease progression, only reaching SCA3 controls performance at week 31 (Figure 4G). Stress exposure for 6 weeks had no impact on muscular strength (Figure 4H). Stressed-WT mice recovered their body weight gain, reaching the levels of non-stressed WT controls 2 weeks after CUS (Figure 4I). Stressed-SCA3 mice took longer to recover their weight gain, being comparable to SCA3 controls only 4 weeks after CUS protocol (Figure 4I).

Assessments of spontaneous horizontal movement and gait showed no differences in SCA3/MJD progression after the application of CUS (Supplementary Figure S2E-F).

## 4. Discussion

Every day we are exposed to stressful events, and our body can effectively adapt and respond, by activating the HPA-axis and consequent glucocorticoid release (reviewed in ^17^). Effective regulation of the stress response is crucial, as improper or extended activation of the HPA axis can be energetically demanding and is associated with various physiological and psychological disorders ^28-31^. In pathological conditions, from major depression to neurodegenerative diseases, the defense mechanisms may be compromised, and the response to stress may be overactivated ^28, 32-35^. In this work, we performed a longitudinal characterization of CORT levels in CMVMJD135 mice during SCA3/MJD progression and explored their ability to recover from stressful events. Stress resilience seems to be a hallmark of this mouse model, as no major differences were observed after acute stress episodes throughout SCA3/MJD progression, and only transient to mild effects were observed after chronic stress exposure in the early stages of the disease. In fact, peripheral CORT levels increase over time in SCA3 mice, showing higher levels in serum at mid to late stages of the disease, starting between 26 and 30 weeks of age.

Regarding our previous results, the reduced GR in the brainstem could justify the elevation of CORT in serum of SCA3 mice ^11^, especially if altered in network regions that interplay with circadian response. However, measurements of CORT receptors, including the mineralocorticoid receptor (MR, which has a higher affinity for CORT ^20, 36^) and GR, at the hypothalamus of our SCA3 mouse are still lacking. This is necessary to further correlate stress response and circadian fluctuations, herein explained by physiological CORT levels, with the role of CORT receptors for feedback regulation of the HPA axis in the SCA3/MJD brain. In fact, de Kloet and colleagues have found that MR blockade but not GR blockade leads to an increased cortisol response to psychosocial stress, in humans ^36^. The authors also highlight that when MR blockade occurs, it results in a lack of fast negative feedback processes, emphasizing the participation of MR during early stress phase responses ^36^. This means that without the proper functioning of MRs, the body’s ability to regulate cortisol release is impaired, leading to prolonged or excessive cortisol levels in response to stress. Nevertheless, in both rats ^37, 38^ and humans ^36^, MR antagonists increased basal and stress-induced cortisol secretion, while GR antagonists had no effect on basal activity but attenuated the initial HPA stress response, leading to prolonged cortisol secretion, due to inhibition of GR negative feedback. This lack of effect of GR blockade on basal HPA-axis activity may be expected as GR is usually not activated by the low glucocorticoid levels secreted during the circadian nadir in basal conditions (reviewed in ^33^). Despite the decrease of GR in the brainstem of SCA3 mice, our present results proved that these mice could functionally regulate HPA-dependent stress responses, herein measured by CORT fluctuations, when exposed to acute stress or HPA-axis suppression events. However, molecular analysis of MR levels and GR overexpression effects still need to be further explored to elucidate the role of corticoid receptors in the SCA3/MJD brain.

While the exact relationship between stress disorders and SCA3 pathogenesis is not well-established, studies on other neurodegenerative, including polyglutamine diseases, have suggested that chronic stress can worsen disease prospects ^35, 39^. On other hand, literature reports that molecular mechanisms like oxidative stress and neuroinflammation may be influenced by chronic stress exposure on neurodegenerative diseases or mood disorders (reviewed in ^40-43^), thus influencing brain homeostasis and pathogenic protein misfolding ^44^. These mechanisms have also been reported to be altered in the context of SCA3/MJD, which may contribute to neuronal dysfunction and death ^4, 45-48^. Our results demonstrate that SCA3 mice under chronic stress exposure showed a worst motor performance upon 6 weeks of CUS when compared to their genotype controls, suggesting an aggravation of their motor phenotype when exposed to stressful events; nevertheless, as disease progressed, stressed SCA3 mice recovered to the levels of SCA3 controls. Therefore, we conclude that chronic stress does not present a major impact on SCA3/MJD progression in mice, with only mild and transient negative effects.

While susceptibility of mice to stressful events seems not to be the case in a SCA3/MJD background, our results indicate that particularly SCA3 female mice have significantly lighter adrenal glands when compared to WT of same sex. In fact, a recent study in mice showed that there is a main sex difference referring to endocrine glands weight, in which female mice had heavier adrenal glands and thymus compared to males ^49^. Nonetheless, the study found that pro-inflammatory consequences of chronic stress exposure were limited, having a similar pattern between sexes in plasma, regardless of stress exposure, whereas the brain exhibited a region-, sex-, and stress-dependent inflammatory pattern ^49^. These findings on female mice adrenal glands weight bring background that could help us justify that the observed genotypic difference for SCA3 females was contributing to the chronic elevation of peripheric CORT at mid-to late-stages of disease. However, this relationship is not supported in male SCA3 mice, as their adrenal glands were not different from WT males and our study was performed using samples from both sexes. Further studies are needed to address possible sex inference on the adrenal glands’ histology in SCA3 mice. Additionally, it may be relevant to instigate endocrine studies at the hypothalamus and anterior pituitary gland level, with quantification of the corticotropin-releasing (CRH) and the adrenocorticotropic (ACTH) hormones in SCA3 mouse brain, as these measures could help us further relate the peripheric elevation of CORT with GR-dependent mechanisms and its impact on SCA3/MJD progression.

Recent advances have revealed the HPA axis as a dynamic homeostatic system, with circadian rhythmicity of glucocorticoid hormones comprising varying amplitude pulses generated in the subhypothalamic nucleus (reviewed in ^33^). This pulsatile output interacts with regulatory networks within the axis, including the adrenal gland, and needs to be precisely decoded at the cellular level. Disrupting this ultradian signal, such as by administering synthetic glucocorticoids, can negatively impact glucocorticoid-dependent systems ^50, 51^, while even subtle alterations in HPA axis dynamics during stress or disease can alter metabolic, behavioral, and cognitive functions ^33, 52, 53^. The development of novel chronotherapies that deliver both circadian and ultradian glucocorticoid patterns holds promise to address this matter. This therapeutic strategy is particularly relevant for progressive neurodegenerative diseases as the timing of treatment may be crucial for targeting specific pathological processes. In the context of SCA3/MJD, little is known about the interplay between the observed neurodegenerative processes and circadian rhythm. However, studying these systems in SCA3/MJD and its molecular redouts pave the way for biomarker breakthroughs that could target specific stages of the disease, thus helping to improve patient progression and follow-up.

## 5. Conclusion

Elevated basal CORT levels and adrenal gland atrophy indicate chronic HPA-axis activation in SCA3 mice, reflecting the systemic impact of the disease. Here we demonstrated that the HPA-axis in SCA3 mice remains responsive to acute stress throughout disease progression, with the ability to restore CORT levels to baseline even at late stages. Thus, the increased CORT levels observed under basal conditions may be related to reduced levels of the glucocorticoid receptor (GR) without altering its intrinsic function. Our findings support our previous hypothesis that the GR decrease is due to its interaction with mutant ATXN3, which may promote its degradation by the proteasome.

Despite chronic stress influencing weight gain and inducing depressive-like behaviors as expected, it does not appear to accelerate the progression of the motor phenotype in SCA3/MJD. Instead, there is an immediate aggravation of animals’ motor performance following stress exposure cessation, which is not sustained as the disease progresses. These insights into the HPA-axis function and stress responses in SCA3/MJD provide a deeper understanding of the disease’s systemic effects.

## Funding

This research was funded by Portuguese national funds, through the Foundation for Science and Technology (FCT) – projects UID/06304/2025 and LA/P/0050/2020 with the DOI numbers 10.54499/UID/06304/2025, and 10.54499/LA/P/0050/2020, respectively. This research was also funded by National Ataxia Foundation (NAF, USA). A. T.-C. received the FCT Exploratory Project (2023.15102.PEX) and the NAF Early Career Investigator Award. FCT funded individual fellowships to J. S. C. (SFRH/BD/140624/2018), D. M.-F. (SFRH/BD/147947/2019), P.G. (PD/BD/135271/2017), B. F.-L. (2024.01049.BD), S. G. (2022.11724.BD), D. C.-G. (2021.08121.BD) and to S. D.-S. (CEECIND/00685/2020).

## The authors declare no conflict of interest

## Supplementary Data

**Figure S1.**
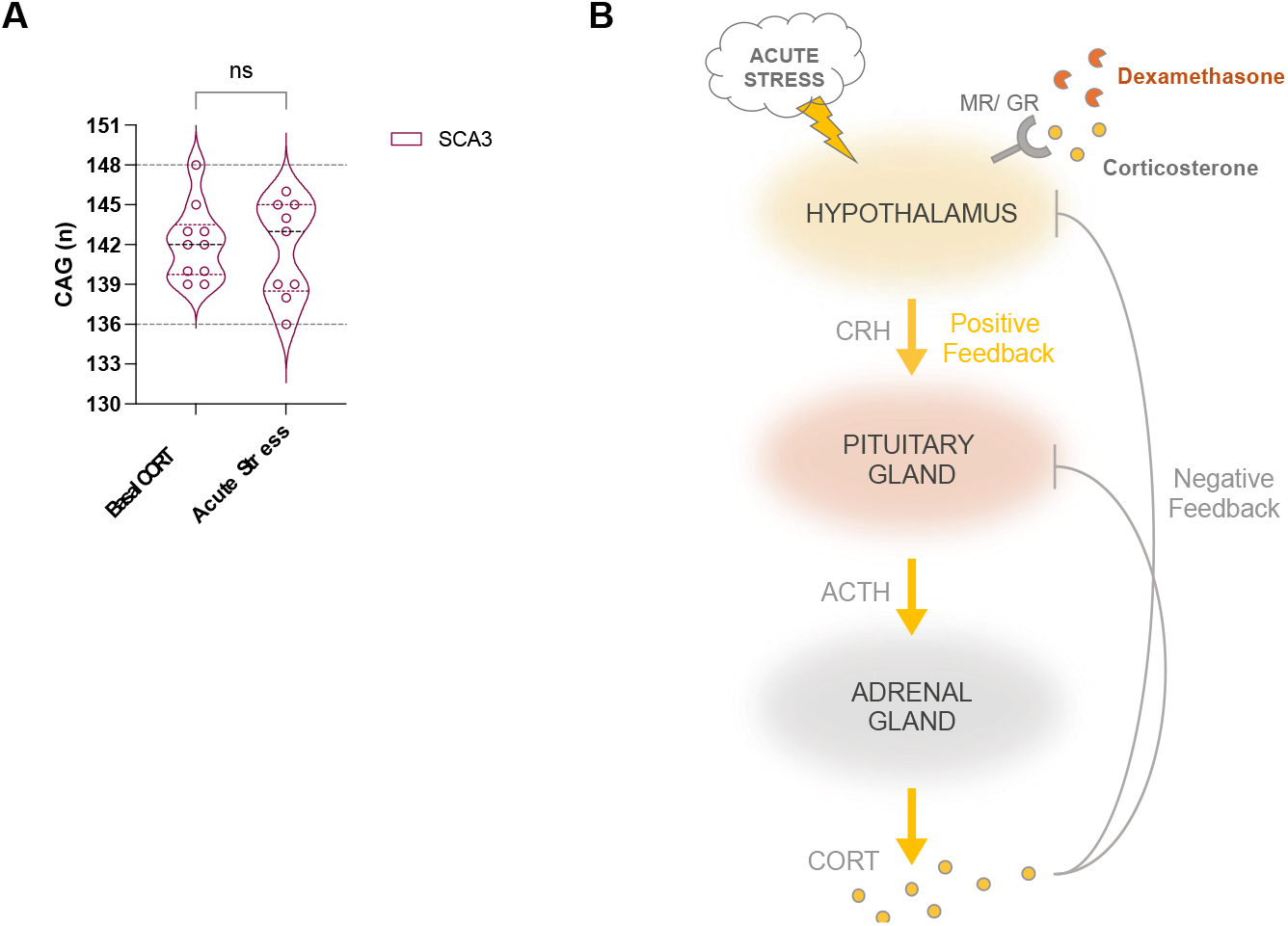
CORT variation assessment groups and HPA-axis explanation. (A) No statistical differences were observed between the CAGn mean of the basal corticosterone (CORT) assessment and the acute stress groups. (B) Schematic representation of the HPA-axis positive feedback upon acute stress stimuli of the hypothalamus and the CORT release cessation at the adrenal gland level through negative-feedback loop dynamics. Both Corticosterone and Dexamethasone compete for the same receptors. Dexamethasone was used to suppress the HPA-axis hormonal response. CAGn comparisons were analyzed using a nonparametric Mann-Whitney U test (A). SCA3 – CMVMJD135 mice; MR – mineralocorticoid receptor; GR – glucocorticoid receptor. Data is represented as the mean ± SEM. ns – not significant.

**Figure S2.**
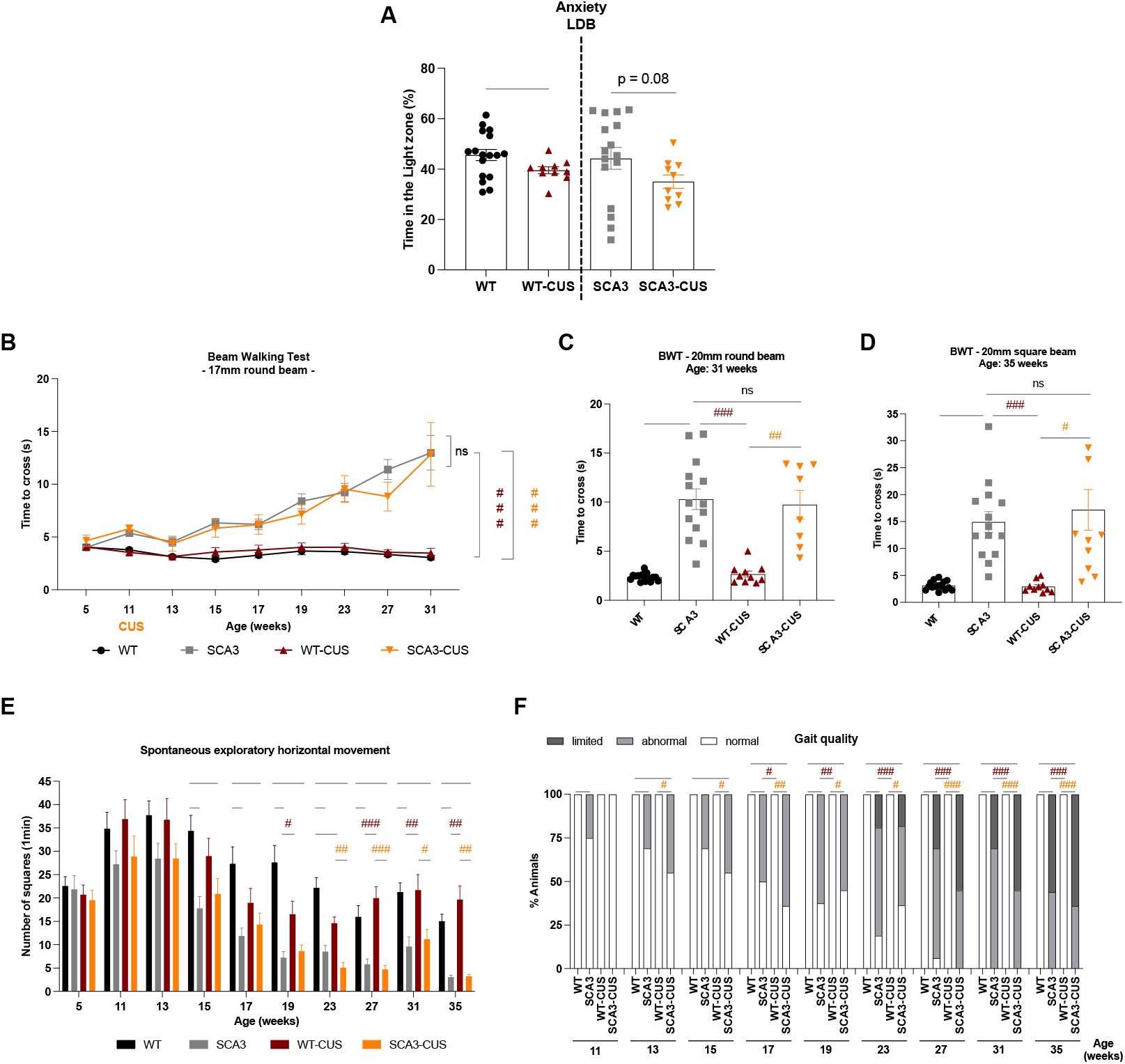
Stressed SCA3 mice showed no differences to control animals in balance, spontaneous movement or gait deficits. (A) A 6-weeks period of CUS could induce anxious phenotype in WT stressed mice (WT-CUS) when compared to WT non-stressed controls, using the light-dark box (LDB) test. No differences in anxiety were observed between SCA3 groups. (B-D) CUS exposure had no impact on disease progression of stressed SCA3 mice (SCA3-CUS) as measured by the time to travel (B) the 17mm round beam between 5 and 31 weeks of age, as well as the (C) 20mm round beam at 31 weeks of age or the (D) 20mm squared beam at 35 weeks of age. (E) Spontaneous horizontal movement and (F) gait quality of SCA3-CUS progressively worsened, but no differences to SCA3 control mice were found. Statistical significance of continuous variables between two groups were analyzed using Student’s t-test to check anxious phenotype between WT and WT-CUS (A – left comparison), and between SCA3 and SCA3-CUS (A - right comparison). A repeated measures ANOVA (B) and two-way ANOVA (C-D) were applied in statistical analyses of continuous variables with normal distribution. Discrete (E) and categorical (F) variables were analyzed using a nonparametric Kruskal-Wallis H test. WT – control wild-type mice; WT-CUS – stressed wild-type mice; SCA3 – control CMVMJD135 mice; SCA3-CUS – stressed CMVMJD135 mice. Data is represented as the mean ± SEM. * represents statistical differences to non-stressed WT group; # represents statistical differences to WT-CUS group; *, # p < 0.05; **, ## p <0.01; ***, ### p <0.001; ns – not significant.

## Notes

### Competing Interest Statement

The authors have declared no competing interest.

