## Supplementary figures and images for "Understanding corticosterone fluctuations and HPA-axis regulation in a mouse model of Spinocerebellar ataxia type 3"

### Supplementary figures

Supplementary figure S1 → fig 2

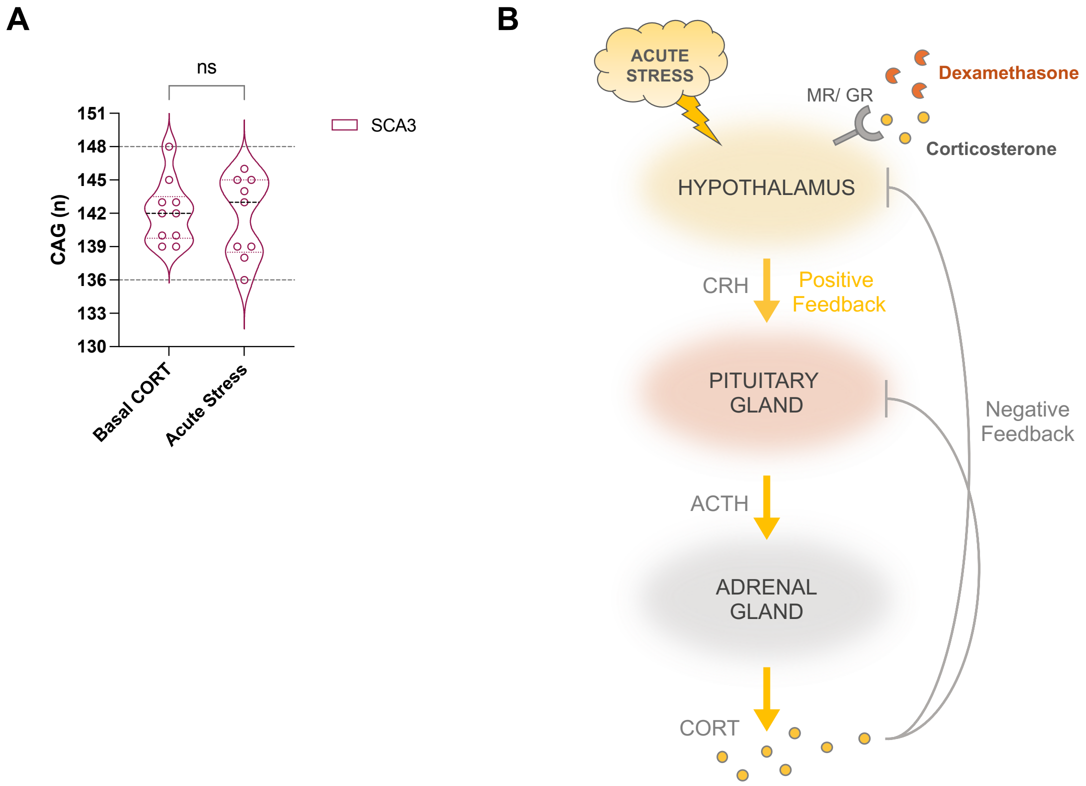

Supplementary figure S2 -→ fig 4

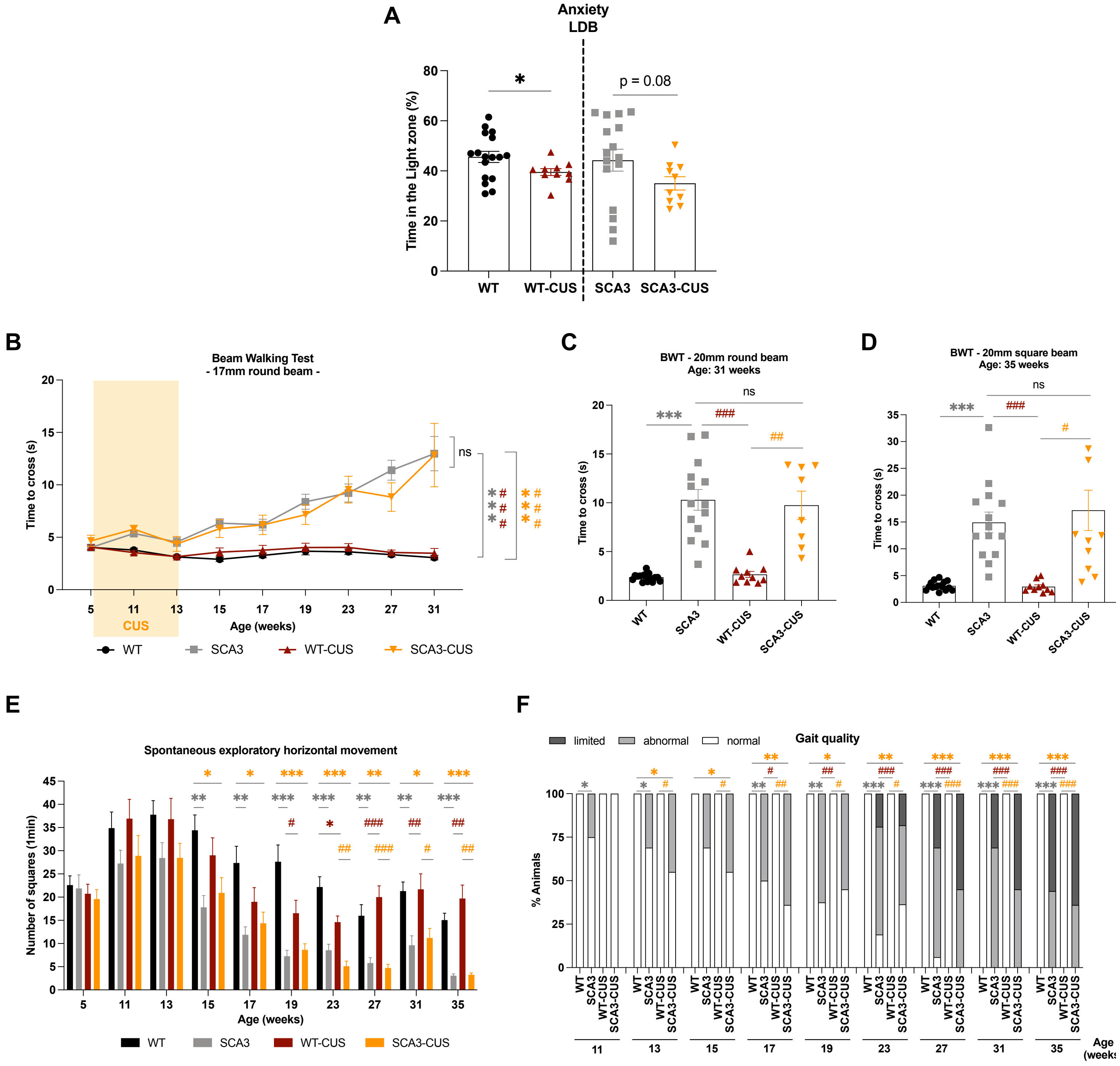
